# Conjugated activation of myocardial-specific transcription of *Gja5* by a pair of Nkx2-5-Shox2 co-responsive elements

**DOI:** 10.1101/2020.05.07.083170

**Authors:** Tianfang Yang, Zhen Huang, Hua Li, Linyan Wang, YiPing Chen

**Author notes:** These authors contributed equally to this work (Tianfang Yang, Zhen Huang).

## Abstract

The sinoatrial node (SAN) is the primary pacemaker in the heart. During cardiogenesis, *Shox2* and *Nkx2-5* are co-expressed in the junction domain of the SAN and regulate pacemaker cell fate through a Shox2-Nkx2-5 antagonism. Cx40 is a marker of working myocardium and an Nkx2-5 transcriptional output antagonized by Shox2, but the underlying regulatory mechanisms remain elusive. Here we characterized a bona fide myocardial-specific *Gja5* (coding gene of Cx40) distal enhancer consisting of a pair of Nkx2-5 and Shox2 co-bound elements in the regulatory region of *Gja5*. Transgenic reporter assays revealed that neither element alone, but the conjugation of both elements together, drives myocardial-specific transcription. Genetic analyses confirmed that the activation of this enhancer depends on Nkx2-5 but is inhibited by Shox2 *in vivo*, and its presence is essential for *Gja5* expression in the myocardium but not the endothelial cells of the heart. Furthermore, chromatin conformation analysis showed an Nkx2-5-dependent loop formation between these two elements and the *Gja5* promoter *in vivo*, indicating that Nkx2-5 bridges the conjugated activation of this enhancer by pairing the two elements to the *Gja5* promoter.

## Introduction

The morphogenesis and physiological maturation of the cardiac conduction system are complicatedly coupled processes, which require a precise spatiotemporal expression of various transcription factors, including Shox2 and Nkx2-5 (Espinoza-Lewis et al., 2009; Hatcher and Basson, 2009; Pashmforoush et al., 2004; van Weerd and Christoffels, 2016). *Nkx2-5* is essential for murine cardiogenesis (Lyons et al., 1995) but its expression was thought to be excluded from the SAN (Espinoza-Lewis et al., 2009), and ectopic expression of Nkx2-5 in the developing SAN inhibits pacemaker cell differentiation (Espinoza-Lewis et al., 2011). In contrast, *Shox2* is essential for SAN development and represses *Nkx2-5* expression in the developing SAN head domain (Espinoza-Lewis et al., 2009; Liu et al., 2014), thus ensuring the absence of Nkx2-5 in the SAN head from the surrounding *Nkx2-5*^+^ working myocardium. It was recently demonstrated the existence of a *Shox2* and *Nkx2-5* co-expression domain in the developing SAN, genetically dividing the SAN into a *Shox2*^+^/*Nkx2-5*^−^ “head” domain and a *Shox2*^+^/*Nkx2-5*^+^ “jFunction” domain (Li et al., 2019; Ye et al., 2015). In the *Shox2*^+^/*Nkx2-5*^+^ SAN junction domain, a Shox2-Nkx2-5 antagonistic mechanism appears to direct the pacemaker cell fate (Ye et al., 2015).

Gap junction alpha-5 (Gja5)/connexin 40 (Cx40), encoded by *Gja5* in mice, plays a critical role in mediating electrical conduction and diffusion of cellular substances in the heart (Bagwe et al., 2005; de Wit et al., 2003; Simon and McWhorter, 2002). Mutations in *GJA5* in humans are associated with atrial fibrillation (AF) (Bai, 2014; Firouzi et al., 2004; Gollob et al., 2006; Juang et al., 2007) and tetralogy of Fallot (TOF) (Greenway et al., 2009; Guida et al., 2013). During murine cardiogenesis, *Gja5* is selectively expressed in the atrial myocardium, the ventricular conduction system, and the arterial endothelial cells (Beyer et al., 2011; Christoffels and Moorman, 2009; van Weerd and Christoffels, 2016). Cardiac *Gja5* expression depends on Nkx2-5 (Dupays et al., 2005; Espinoza-Lewis et al., 2011; Linhares et al., 2004) but is repressed by Shox2 (Blaschke et al., 2007; Espinoza-Lewis et al., 2009), and is controlled tightly by the Shox2-Nkx2-5 antagonism in the *Nkx2-5*^+^/*Shox2*^+^ domains (Li et al., 2019; Ye et al., 2015). However, the underlying mechanisms utilized by Shox2 and Nkx2-5 to control *Gja5* expression remain unknown.

The precise spatiotemporal control of transcriptional networks is critical for cardiac development (Bruneau, 2008; Olson, 2006). Whereas proximal promoters provide a basis for gene transcription, the tissue-specific gene expression is dependent mainly on the interactions between existing tissue-specific transcription factors and a variety of regulatory DNA sequences classified as distal enhancers (Levine, 2010; Nord et al., 2013; Visel et al., 2009). In the heart, numerous cardiac-specific distal enhancers have been characterized by comparative genomic analyses searching for evolutionarily conserved elements and genome-wide profiling of active enhancer associated epigenetic marks or co-activator protein binding sites in heart tissues (Blow et al., 2010; Dickel et al., 2016; He et al., 2011; May et al., 2012; Narlikar et al., 2010; Paige et al., 2012; Wamstad et al., 2012). Although the combinatorial binding of cardiac transcription factors at proximal promoters and their physiological significance in transcription regulation have been studied extensively (Bruneau, 2002), the functions of cardiac-specific distal enhancers and how transcription factors regulate them remain largely unexplored.

In this study, we report the characterization of a myocardial-specific *Gja5* distal enhancer, which is formed by a pair of Shox2 and Nkx2-5 co-bound elements (named as *Gja5-S1* and *Gja5-S2*) spaced by a 12-Kb non-coding region downstream of *Gja5*. Neither *Gja5-S1* nor *Gja5-S2* alone, but the conjugation of both elements termed as *Gja5-eh*, displayed robust enhancer activity recapitulating the endogenous *Gja5* expression pattern in the *Nkx2-5*^+^ domain of the developing and adult heart. Genetic analyses showed that the *Gja5-eh* enhancer activity is indeed dependent on Nkx2-5 but is inhibited by Shox2 *in vivo*. Mice bearing the ablation of both *Gja5-S1* and *Gja5-S2* exhibit drastically reduced endogenous *Gja5* expression only in the myocardium but not the arterial endothelial cells, illustrating that these two elements are indispensable for *Gja5* expression specifically in the myocardium *in vivo*. Moreover, chromatin conformation analysis of mouse embryonic hearts showed that both *Gja5-S1* and *Gja5-S2* have frequent contacts with the *Gja5* promoter in an Nkx2-5-dependent manner, indicating that Nkx2-5 bridges the enhancer activation by conjugating *Gja5-S1* and *Gja5-S2* together with the *Gja5* promoter.

## Results and Discussion

### Identification of *Gja5-S1* and *Gja5-S2* and generation of *Gja5-eh-LacZ* reporter mouse line

To understand the functional mechanisms underlying the Shox2-Nkx2-5 antagonism that regulates cell fate in the developing SAN, we revisited the genomic profiles of Shox2 and Nkx2-5 ChIP-seq assays on the right atrial tissues of E12.5 mouse embryonic heart, which displayed a substantial genome-wide co-occupancy of Shox2 and Nkx2-5 (Ye et al., 2015). Among the co-binding peaks, two sites (termed *Gja5-S1* and *Gja5-S2*) co-occupied by Shox2 and Nkx2-5 downstream of *Gja5* raised our particular interest (Fig. 1A), because *Gja5* expression has been proven to be regulated by *Shox2* and *Nkx2-5* during SAN development (Li et al., 2019; Ye et al., 2015). *Gja5-S1* locates at around 9-Kb downstream of the *Gja5* coding region, whereas *Gja5-S2* sits downstream of *Gja5-S1* separated by a 12-Kb non-coding sequence. Additional analysis integrating accessible public data showed that *Gja5-S1* and *Gja5-S2* are located in the same topological associated domain (TAD) as *Gja5* (Dixon et al., 2012), and are both marked as putative open chromatin in embryonic hearts but not in any other tissues at the same stage (ENCODE Project Consortium, 2012) (Fig. S1), further suggesting that these two elements may have cardiac-specific enhancer activity.

**Figure 1.**
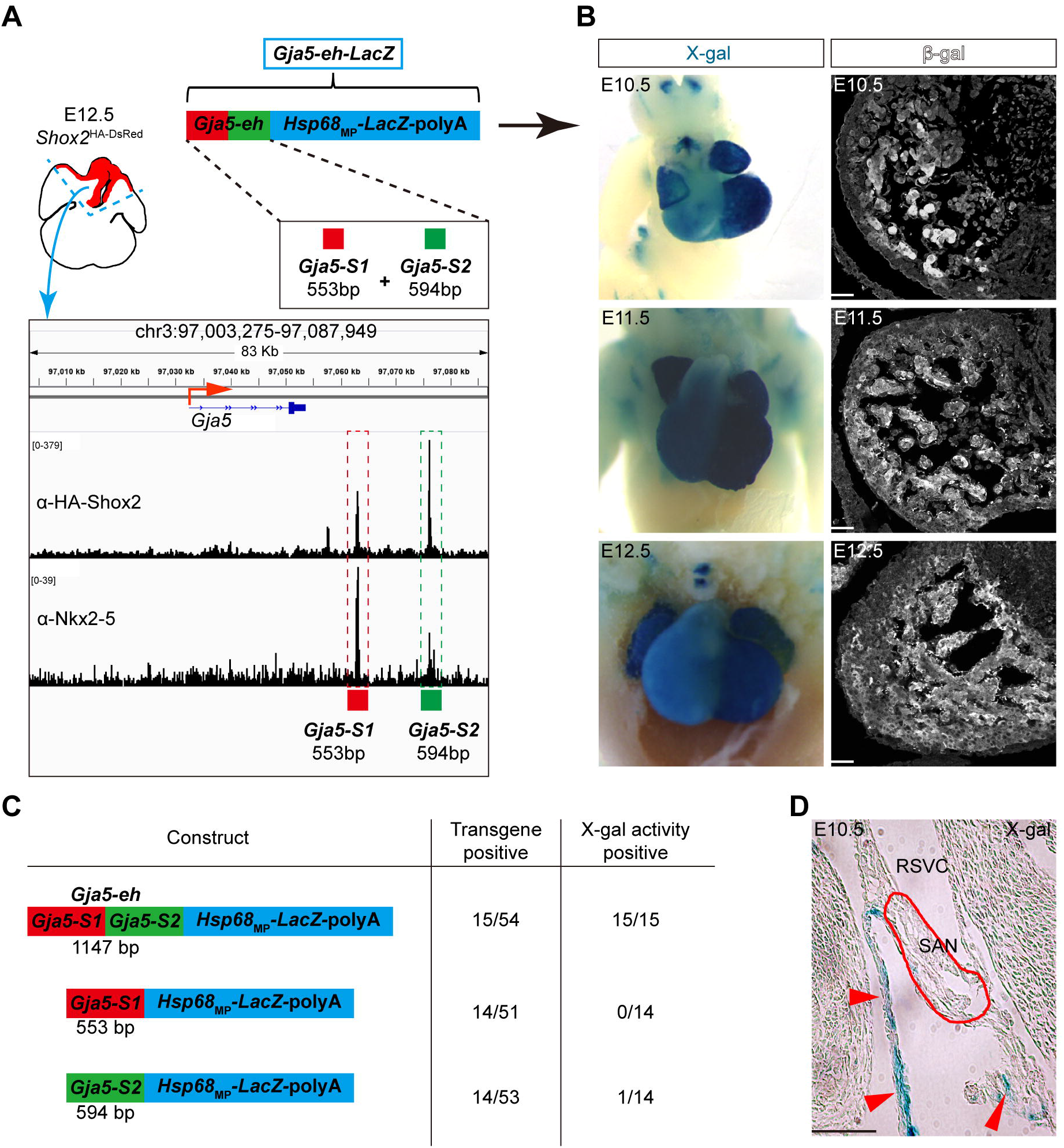
Identification of *Gja5-S1* and *Gja5-S2* and generation of *Gja5-eh-LacZ* reporter mice. (A) Schematic overview of ChIP-seq data collection, visualization, and identification of *Gja5-S1* and *Gja5-S2*, and generation of *Gja5-eh-LacZ* transgenic construct. (B) Whole-mount X-gal staining and section immunostaining of β-galactosidase of E10.5, E11.5 and E12.5 *Gja5-eh-LacZ* embryos. (C) Schematic diagram summarizing the reporter constructs of *Gja5-S1*, *Gja5-S2*, and *Gja5-eh*. The left column indicates the design of each construct. The middle column indicates the number of transgene-positive F0 embryos as a fraction of the total number of F0 embryos. The right column indicates the number of F0 embryos showing cardiac-specific X-gal activity as a fraction of the total number of transgene-positive F0 embryos. (D) X-gal staining on a cryo-sectioned E10.5 *Gja5-eh-LacZ* embryo. Red arrowheads point to positive X-gal staining in the atrial tissue. Note that the staining is excluded from the SAN (circled in red). RSVC, right superior vena cava. SAN, sinoatrial node. Scale bars: 50 μm.

To assess the enhancer activity of *Gja5-S1* and *Gja5-S2 in vivo*, we initially cloned each of them into a reporter construct containing the *Hsp68* minimal promoter (*Hsp68*_mp_) and β-galactosidase coding sequence (*LacZ*) (Kothary et al., 1989). However, neither element exhibited enhancer activity in transient transgenic reporter assays (Fig. 1C). We wondered whether the two elements are required to function together, and therefore subsequently conjugated both elements together as a fusion fragment, termed *Gja5-eh*, and generated a reporter construct named *Gja5-eh-LacZ* to test its enhancer activity in transgenic mouse embryos. Strikingly, all the *Gja5-eh-LacZ* transgene-positive embryos (15/15) showed beta-galactosidase (β-gal) activity in the developing heart (Fig. 1B, C). Section immunostaining and X-gal staining further revealed the specific expression of *LacZ* in the myocardium but its absence in the SAN of the transgenic animals, similar to the pattern of endogenous *Gja5* expression (Fig. 1B, D). These observations indicate that both *Gja5-S1* and *Gja5-S2* are required to act together *in cis* to drive cardiac-specific gene transcription. We subsequently generated *Gja5-eh-LacZ* permanent transgenic reporter mouse lines to facilitate further studies.

### *Gja5-eh* recapitulates endogenous *Gja5* expression in *Nkx2-5*^+^ myocardium

Next, we examined whether the enhancer activity of *Gja5-eh* could faithfully recapitulate the endogenous *Gja5* expression in the developing heart. Since Nkx2-5 binds to *Gja5-S1* and *Gja5-S2* and is essential for *Gja5* expression in the heart (Dupays et al., 2005), we conducted co-immunostaining on Nkx2-5, Cx40, and β-gal in the hearts of *Gja5-eh-LacZ* reporter mice. The results showed co-expression of β-gal and Cx40 in the *Nkx2-5*^+^ myocardium of the ventricle and atrium of *Gja5-eh-LacZ* hearts, not only at the embryonic stage (E12.5) but also at adulthood (P60) (Fig. 2A-D, single-channel images in Fig. S2). Similar to the results seen in the transient transgenic reporter assay, β-gal was not detectable in the *Shox2*^+^ SAN region in contrast to its presence in the adjacent Nkx2-5^+^/Cx40^+^ atrial myocardium (Fig. 2A, Fig. S2M-P). Notably, we also observed the absence of β-gal in the Nkx2-5^−^/Cx40^+^ coronary arteries (CA) in contrast to its intense expression in the Nkx2-5^+^/Cx40^+^ ventricular trabeculae (VT) (Fig. 2D, Fig. S2I-L). Similar observations were found in other developmental stages (data not shown). These results demonstrate that *Gja5-eh* acts as a functional distal enhancer and recapitulates the endogenous *Gja5* expression. However, its enhancer activity is restricted only to the Nkx2-5^+^ myocardium, suggesting a positive role of Nkx2-5 in the activation of the enhancer.

**Figure 2.**
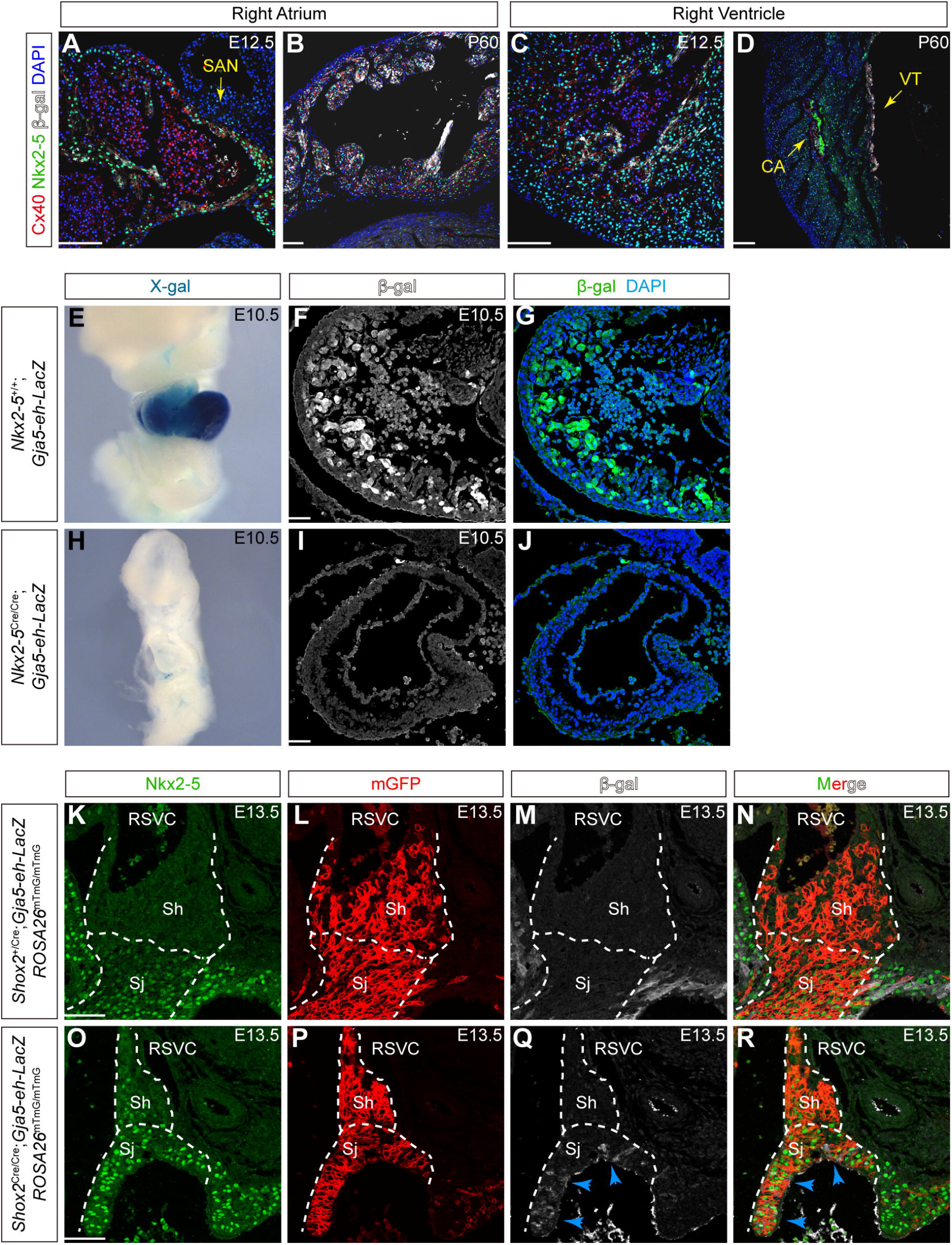
*Gja5-eh* enhancer activity recapitulates endogenous *Gja5* expression in *Nkx2-5*^+^ myocardium, and is dependent on Nkx2-5 but is inhibited by Shox2. (A-D) Triple immunofluorescent staining (Nkx2-5, Cx40, β-gal) on right atrium (A,B) and right ventricle (C,D) of the *Gja5-eh-LacZ* hearts at E12.5 and P60. CA, coronary arteries; VT, ventricular trabeculae; SAN, sinoatrial node. Scale bars: 100 μm. Also see Fig. S2. (E-J) *Gja5-eh-LacZ* transgenic mice were crossed onto *Nkx2-5*^+/+^ (E-G) and *Nkx2-5*^Cre/Cre^ (H-J) backgrounds and enhancer activity was examined by whole-mount X-gal staining and fluorescent immunostaining for β-galactosidase at E10.5. Scale bars: 50 μm. (K-R) *Gja5-eh-LacZ* transgenic mice were crossed onto *Shox2*^+/Cre^;*ROSA26*^mTmG/mTmG^ (K-N) and *Shox2*^Cre/Cre^;*ROSA26*^mTmG/mTmG^ (O-R) backgrounds, and examined at E13.5 for expression of Nkx2-5, mGFP and β-galactosidase by triple fluorescent immunostaining. Blue arrowheads point to ectopic β-gal signal co-localized with Nkx2-5 and mGFP in the SAN junction domain. RSVC, right superior vena cava; Sh, SAN head; Sj, SAN junction. Scale bars: 50 μm.

### *Gja5-eh* enhancer activity is dependent on Nkx2-5 but is inhibited by Shox2

To determine whether Nkx2-5 regulates *Gja5-eh* enhancer activity *in vivo*, we compounded *Gja5-eh-LacZ* allele onto an *Nkx2-5*-null background using the *Nkx2-5*^Cre^ knock-in allele (Moses et al., 2001; Ye et al., 2015). As we expected, the enhancer activity of *Gja5-eh* was abolished in the *Nkx2-5*^Cre/Cre^;*Gja5-eh-LacZ* embryonic hearts as compared to controls (Fig. 2E-J), indicating an indispensable role of Nkx2-5 in the activation of *Gja5-eh*. Since Shox2 also binds to *Gja5-S1* and *Gja5-S2* while represses *Gja5* expression in the heart (Espinoza-Lewis et al., 2009; Ye et al., 2015), we next examined whether Shox2 inhibits *Gja5-eh* enhancer activity *in vivo*. To do this, we compounded *Gja5-eh-LacZ* allele with *ROSA26*^mTmG^ and *Shox2*^Cre^ alleles to generate *Shox2*^Cre/Cre^;*Gja5-eh-LacZ*;*ROSA26*^mTmG^ mice, which enabled monitoring of *Gja5-eh* enhancer activity and tracing of *Shox2* lineage in *Shox2*-null background (Sun et al., 2013). In contrast to the absence of β-gal in the *Nkx2-5*^+^ cells derived from the *Shox2* lineage cells that were labeled by mGFP in controls (*Shox2*^+/Cre^;*Gja5-eh-LacZ*;*ROSA26*^mTmG^ (Fig. 2K-N), we observed the presence of *Nkx2-5*^+^/mGFP^+^/β-gal^+^ cells in the SAN junction, but not the SAN head domain, of *Shox2*^Cre/Cre^;*Gja5-eh-LacZ*;*ROSA26*^mTmG^ mice (Fig. 2O-R), indicating the cell-autonomous ectopic activation of *Gja5-eh* enhancer in the absence of *Shox2*. This result is consistent with the ectopic expression of *Gja5* in the cardiac structures in *Shox2*-null embryos (Espinoza-Lewis et al., 2009; Sun et al., 2015), suggesting that Shox2 suppresses *Gja5* expression by binding to *Gja5-S1* and *Gja5-S2*.

### *Gja5-S1* and *Gja5-S2* are essential for myocardial *Gja5* expression

To investigate whether *Gja5-S1* and *Gja5-S2* serve as a bona fide distal enhancer that drives endogenous *Gja5* expression, we generated a knockout allele termed *Gja5*^Δ14^, which carries a deletion of a 14-Kb region that includes both *Gja5-S1*, *Gja5-S2*, and the sequence between these two sites (Fig. 3A). As a control, we also generated a pseudo-knockout allele termed *Gja5*^Δ12^, in which the 12-Kb sequence between *Gja5-S1* and *Gja5-S2* was deleted (Fig. 3B). We subsequently examined and compared the endogenous *Gja5* expression in the hearts of *Gja5*^Δ14/Δ14^, *Gja5*^Δ12/Δ12^, as well as wild type mice. Strikingly, we observed that in the perinatal *Gja5*^Δ14/Δ14^ hearts, Cx40 expression in the atrial and ventricular trabecular myocardium was almost completely abolished but remained unaltered in the coronary arteries (Fig. 3D, G). We also verified by Western blotting the dramatically decreased levels of Cx40 proteins in the atria and ventricles of P0 *Gja5*^Δ14/Δ14^ mice as compared to *Gja5*^+/+^ controls (Fig. 3I). Further quantification via real-time qPCR (RT-qPCR) showed that the *Gja5* transcription levels remained only 5%±0.2% in the atria and 31%±3.7% in the ventricles of *Gja5*^Δ14/Δ14^ mice as compared to their counterparts in *Gja5*^+/+^ controls (Fig. 3J). These residual *Gja5* expressions in *Gja5*^Δ14/Δ14^ groups may largely be contributed by coronary arteries where *Gja5* expression is unaffected. In contrast, immunohistochemical staining revealed indiscernible change in Cx40 expression in both myocardium and coronary arteries in *Gja5*^Δ12/Δ12^ perinatal hearts as compared to that of *Gja5*^+/+^ mice (Fig. 3E, H). These results demonstrate that *Gja5-S1* and *Gja5-S2* are explicitly required in the myocardium to drive *Gja5* expression, consistent with the specific enhancer activity of *Gja5-eh* in the *Nkx2-5*^+^ myocardium, but not in the *Nkx2-5*^−^ coronary arteries (Fig. S2I-L). Despite the diminished myocardial *Gja5* expression in *Gja5*^Δ14/Δ14^ hearts, these mice are viable and fertile with no overt phenotype. We subsequently performed surface ECG on adult *Gja5*^Δ14/Δ14^ mice along with controls but did not find obvious differences between mutants and controls (N = 10 for each genotype, with 5 males and 5 females, data not shown). Indeed, *Gja5* null mutant mice were viable and fertile, with only minor cardiac physiological abnormalities (prolonged P-wave manifested by ECG) (Bagwe et al., 2005). However, simultaneous loss of *Cx37* and *Gja5*(*Cx40*) results in perinatal lethality with severe vascular defect, indicating that *Gja5* functions importantly but redundantly in the endothelial lineage (Simon and McWhorter, 2002). Therefore, the normal cardiac function of *Gja5*^Δ14/Δ14^ mice, revealed by ECG, can be explained by the unaffected expression of *Gja5* in the endothelial lineage. Additionally, although a number of mutations within *GJA5* coding regions have been associated with atrial fibrillation, none was reported occurring in the non-coding regulatory element associated with *GJA5* so far. We speculate that the pathogenicity of mutations in *GJA5* may arise from alterations of biochemical properties of the CX40 protein rather than changes in overall CX40 levels. Admittedly, we acknowledge that the physiological consequences of the 14Kb deletion could be tested in more detail by additional methods, such as echocardiography and optical mapping.

**Figure 3.**
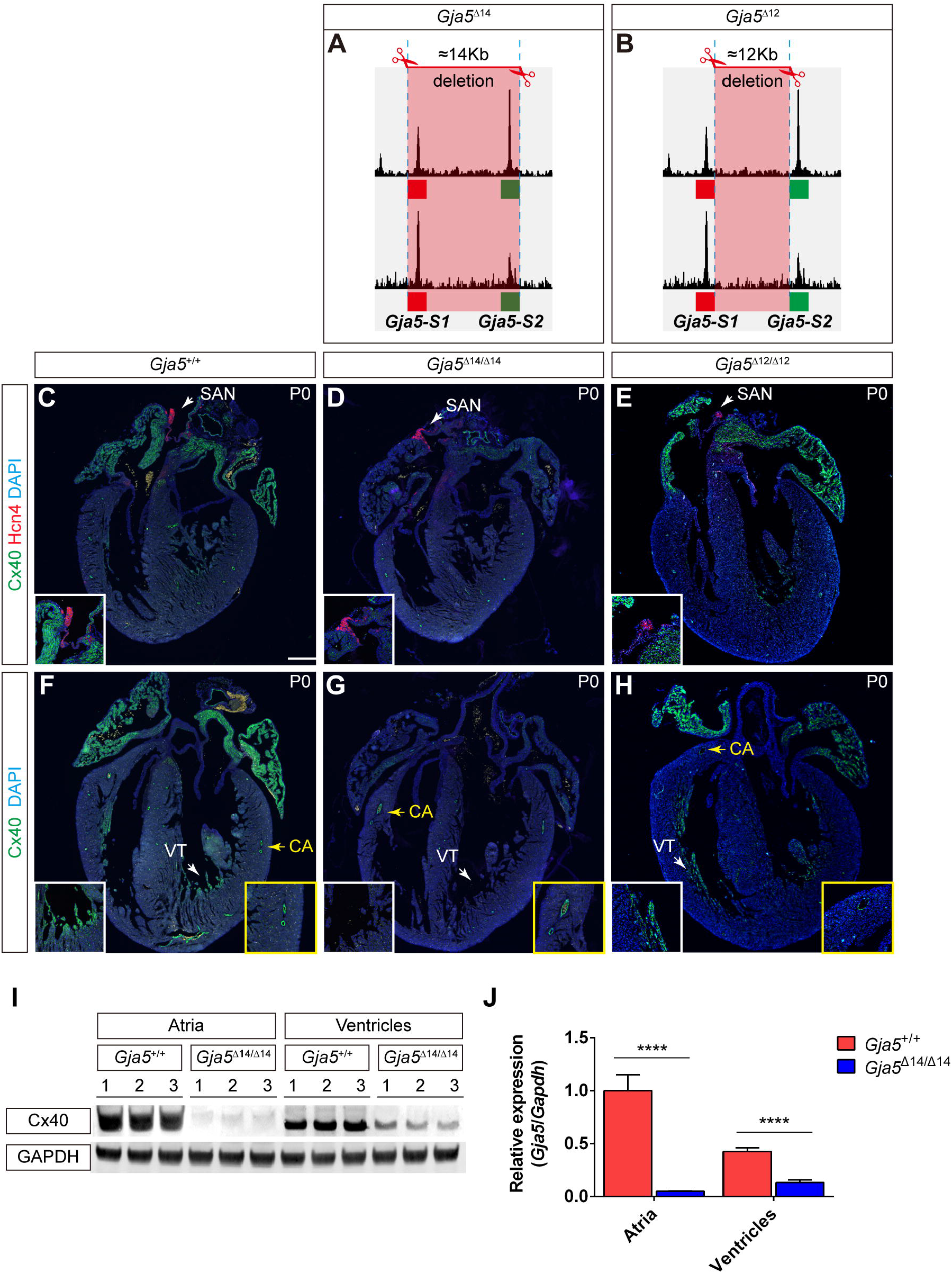
*Gja5-S1* and *Gja5-S2* are essential for myocardial *Gja5* expression. (A, B) Schematic overview of CRISPR/Cas9-mediated mouse genome editing and generation of *Gja5*^Δ14^ and *Gja5*^Δ12^ alleles. (C-H) Double fluorescent immunostaining of Cx40 and Hcn4 on *Gja5*^+/+^ (C, F), *Gja5*^Δ14/Δ14^ (D, G), and *Gja5*^Δ12/Δ12^ (E, H) hearts at P0. White/yellow arrowheads pointed regions are highlighted in white/yellow outlined squares as inserts. SAN, sinoatrial node; VT, ventricular trabeculae; CA, coronary arteries. Scale bars: 250 μm. (I) Western blot of Cx40 and GAPDH on protein extracts from atria or ventricles of *Gja5*^+/+^ and *Gja5*^Δ14/Δ14^ hearts at P0. (J) RT-qPCR detecting *Gja5* and *Gapdh* expression on atria or ventricles of *Gja5*^+/+^ and *Gja5*^Δ14/Δ14^ hearts at P0. Maximum *Gja5* expression relative to *Gapdh* is set as one. ****: P < 0.0001. Biological replicates N = 3.

### An Nkx2-5-dependent interaction between *Gja5-S1*/*Gja5-S2* and the *Gja5* promoter

Recent studies have reported prevalent long-range enhancer-promoter interactions in cardiomyocytes (Montefiori et al., 2018; Rosa-Garrido et al., 2017) and association of disrupted enhancer-promoter looping with cardiac malfunctions (Man et al., 2019). Nkx2-5 was reported to bind to the *Gja5* promoter and upregulate its expression *in vitro* (Linhares et al., 2004), but its functional mechanisms remain elusive. To determine whether *Gja5-S1* and *Gja5-S2* have direct interaction with the *Gja5* promoter and if such interaction requires the presence of Nkx2-5, we performed Chromatin Conformation Capture (3C) (Dekker et al., 2002) assays on E10.5 wild type and *Nkx2-5*^Cre/Cre^ hearts. The results showed that in wild type groups, fragments embracing primer pairs F2+R1 and F4+R1 bore significantly higher relative interaction frequency (RIF) than other primer pairs (Fig. 4A, B), indicating that both *Gja5-S1* and *Gja5-S2* form a strong loop with the *Gja5* promoter. In comparison, the RIF between *Gja5-S1/Gja5-S2* and the *Gja5* promoter of *Nkx2-5*^Cre/Cre^ (*Nkx2-5* null) hearts was significantly reduced (Fig. 4A, B). These results demonstrate an Nkx2-5-dependent loop formation between both *Gja5-S1*/*Gja5-S2* and the *Gja5* promoter, illustrating a critical enhancer-pairing and promoter-docking function of Nkx2-5 in the conjugated activation of *Gja5-S1* and *Gja5-S2*. This bridging function of Nkx2-5 may result from its ability to form homodimers or heterodimerize with other Nkx2 family transcription factors (Kasahara et al., 2001). Interestingly, we did not observe the binding peaks of Nkx2-5 to the *Gja5* promoter from our ChIP-seq data (Fig. 1A, Fig. S1), raising the possibility that Nkx2-5 may have different binding preferences at a context-specific manner.

**Figure 4.**
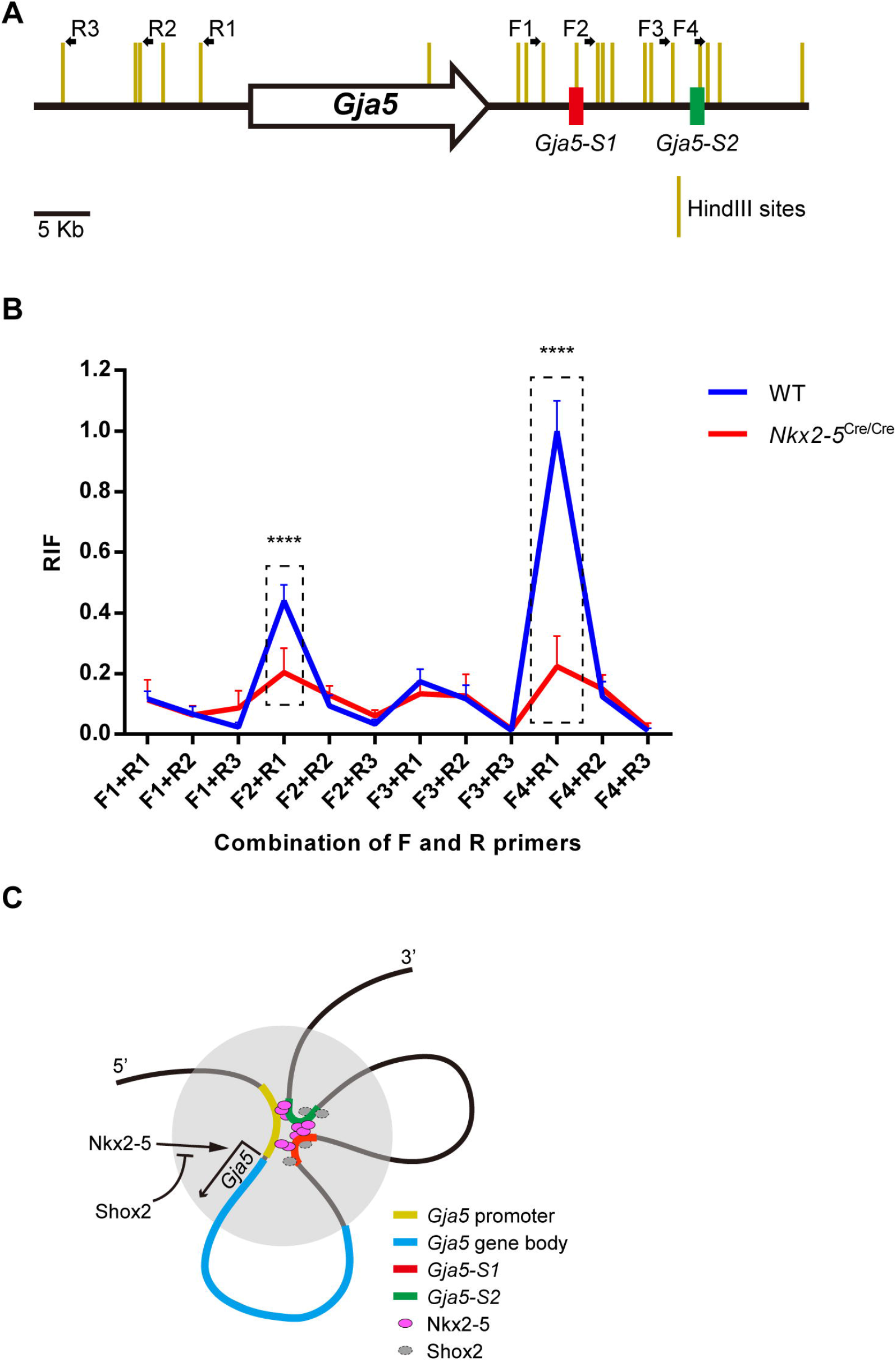
Nkx2-5 mediates the interaction between *Gja5-S1*/*Gja5-S2* and the *Gja5* promoter. (A) Relative positions of primer sets used for 3C assays. (B) Relative interaction frequency (RIF) in 3C assay between a combination of forward and reverse primer sets in E10.5 hearts of wild type and *Nkx2-5*^Cre/Cre^ mice. Maximum RIF is set as one and relative fold changes are shown. ****: P < 0.0001. Biological replicates N = 3. (C) a schematic model proposing that *Gja5-S1* and *Gja5-S2* act in conjugation as a myocardial-specific *Gja5* distal enhancer by interacting with the *Gja5* promoter, and the enhancer activity depends on Nkx2-5 but inhibited by Shox2.

We also looked for consensus binding sites of Nkx2-5 and Shox2 within both *Gja5-S1* and *Gja5-S2* by searching JASPAR-2020, a recently updated database of in vitro DNA binding motifs (Fornes et al., 2019). We found numerous predicted transcription factor binding sites within *Gja5-S1* (Fig. S4) and *Gja5-S2* (Fig. S5). However, there is only one Nkx2-5 binding site (-AACCACTCCAG-) inside *Gja5-S1* but not *Gja5-S2*, while no Shox2 binding site was found. Meanwhile, binding sites of two other NK family transcription factors, Nkx2-3 and Nkx2-8, were found in both *Gja5-S1* (-GCACTTGAG-) and *Gja5-S2* (-CCACTTGAC-). All these three sites share motif -(C/G)CACT(T/C)-, consistent with our Nkx2-5 ChIP-seq results (Ye et al., 2015) and results from another study (He et al., 2011). Our previous studies showed that Shox2 has distinct binding sites among different tissues during organogenesis (Wang et al., 2020; Xu et al., 2019; Ye et al., 2016, 2015), but a common motif is yet to be characterized. The binding preferences of Shox2 may be partially dependent upon other co-factors, such as Hox family transcription factors, as reported previously (Ye et al., 2016).

Based on the work presented here, we propose that *Gja5-S1* and *Gja5-S2* are a pair of Nkx2-5-Shox2 co-responsive elements that act together as a myocardial-specific *Gja5* distal enhancer by interacting with the *Gja5* promoter (Fig. 4C). We acknowledge that as the ChIP-seq and 3C experiments were performed on bulk tissues consisting of both myocardium and SAN, we can not conclude that *Gja5-S1* and *Gja5-S2* are co-bound by both Nkx2-5 and Shox2 simultaneously. One interpretation of the model could be that in the SAN junction, Shox2 competes with Nkx2-5 for binding to these two elements and thus disrupts the interaction of the enhancers with the *Gja5* promoter by keeping the enhancers at a distal status. Another interpretation is that Shox2 and Nkx2-5 form non-functional protein complexes that can not bind to DNA or have diminished DNA binding specificities, since Shox2 and Nkx2-5 have physical interaction *in vitro* (Ye et al., 2015).

While we report that *Gja5-S1* alone does not have enhancer activity (Fig. 1C), a 705-bp *Gja5* enhancer that embraces *Gja5-S1* was reported (Hashimoto et al., 2019). *Gja5-S1* (553-bp) locates in the center of this 705-bp enhancer, flanked by an additional 152-bp sequence (90 bp upstream/62 bp downstream). This additional 152-bp sequence, which contains Hand2 binding sites and is enriched in H3K27ac, appears to contribute to the discrepant enhancer activity via H3K27ac deposition recruited by Hand2 (Creyghton et al., 2010; Hashimoto et al., 2019). In contrast, we found that *Gja5-S2* is marked by H3K4me2, H3K4me3, and occupied by CCCTC-binding factor (CTCF) (Fig. S3) (ENCODE Project Consortium, 2012). Both H3K4me2 and H3K4me3 are associated with active transcription (Heintzman et al., 2007; Kim and Buratowski, 2009; Mikkelsen et al., 2007; Santos-Rosa et al., 2002), and CTCF is a critical regulator of enhancer-promoter interactions and chromatin topology (Barrington et al., 2019; Nora et al., 2017; Shin, 2019; Wutz et al., 2017). These distinct histone modifications and binding of chromatin modifiers between *Gja5-S1* and *Gja5-S2* may display complementary actions that contribute to their conjugated activation. Based on the proposed model, we also speculate that a smaller deletion of either *Gja5-S1* or *Gja5-S2* should abolish *Gja5* expression in the myocardium and phenocopy *Gja5*^Δ14/Δ14^ mice.

The behaviors of enhancers are multifaceted. During organogenesis, tissue-specific enhancers of the same gene could function redundantly to achieve phenotypic robustness by buffering the loss-of-function mutations of individual enhancers (Frankel et al., 2010; Osterwalder et al., 2018). Strong enhancers may compete for contacting promoters, and enhancers of weak or intermediate strength by themselves may function additively to enhance the expression of the same gene (Bothma et al., 2015). One enhancer may contain multiple binding sites for different transcription factors and is synergistically activated through proximal protein-protein interactions (Ambrosetti et al., 1997; Anderson et al., 2017; Grieder et al., 1997; Hashimoto et al., 2019). Here, our studies show that the nature of enhancers could be even more complicated. This conjugated activation of myocardial-specific activation of *Gja5* by two regulatory elements may represent a novel mechanism for accuracy assurance of tissue-specific gene expression. Although *Gja5-eh* is a recombinant fusion of *Gja5-S1* and *Gja5-S2,* we indeed observed a striking consistency between *Gja5-eh-LacZ* enhancer activity and molecular phenotype of *Gja5*^Δ14/Δ14^ mice. We reason that the fusion of *Gja5-S1* and *Gja5-S2* has mimicked their actual close contact at the *Gja5* promoter loci *in vivo*, which makes *Gja5-eh* represent the spatiotemporal behaviors of *Gja5-S1* and *Gja5-S2* precisely. The *Gja5*^Δ12^ allele actually also simulates the fusion of *Gja5-S1* and *Gja5-S2 in situ* and represents their conjugated activation (Fig. 3B, E, H). Collectively, these observations point out a unique pattern of conjugated enhancer activation and provide novel insights into characterization, design, and optimization of tissue/lineage-specific enhancers that may benefit biomedical research or therapeutic applications.

## Materials and Methods

### Cloning and plasmids

The 553-bp *Gja5-S1* and 594-bp *Gja5-S2* fragments were amplified by PCR, respectively, using primers Gja5-S1-F (5’ -AGTCTGATGACAACTTGTGAGAATCG-3’), Gja5-S1-R (5’-GGGTGACAGTAAGAAATGTCAGGTG-3’), and Gja5-S2-F (5’ -AATGAACAGGAAAGTGGGAG G-3’), Gja5-S2-R (5’-CAGGGCGGTCAGGCAG-3’). The 1147 bp fusion fragment *Gja5-eh* was synthesized commercially (Qinglanbiotech.com). Each of the fragments was subsequently cloned into plasmid *hsp68-LacZ* (Kothary et al., 1989) to generate *Gja5-S1-LacZ*, *Gja5-S2-LacZ*, and *Gja5-eh-LacZ* constructs for transient transgenic reporter assays. *Gja5-eh-LacZ* was also used for generating permanent transgenic reporter mouse lines.

### Mouse models

The generation and genotyping methods of *Shox2*^Cre^, *Nkx2-5*^Cre^ and *ROSA26*^mTmG^ mice have been described previously (Moses et al., 2001; Muzumdar et al., 2007; Sun et al., 2013). Generation of transgenic embryos and mice were performed as described previously (Ye et al., 2016). The *Gja5*^Δ14^ and *Gja5*^Δ12^ alleles were generated by CRISPR/Cas9-mediated genome editing (Wang et al., 2013). Additional details of CRISPR/Cas9-mediated genome editing and transgenesis methods are described in the Supplementary Materials and Methods. All animal work in this study was approved by The Tulane University Institutional Animal Care and Use Committee (IACUC). Sample sizes were empirically determined based on previous experimental procedures (Ye et al., 2016). Mouse embryos were excluded from further analysis only if they did not carry alleles of interest.

### Histology, immunohistochemistry, and X-gal staining

Hearts were harvested from properly euthanized mice or staged embryos, fixed in ice-cold 4% paraformaldehyde (PFA) overnight at 4◻, dehydrated through gradient ethanol, cleared in xylene, embedded in paraffin, and sectioned at 5μm for immunostaining as described previously (Li et al., 2019). The primary antibodies used in this study were: anti-Hcn4 (ab32675, Abcam; 1:200), anti-Nkx2-5 (AF2444, Novus Biologicals; 1:200), anti-Cx40 (Cx40-A; Alpha Diagnostic International; 1:200), anti-GFP (sc-9996, Santa Cruz Biotechnology; 1:200), anti-β-galactosidase (ab9361, Abcam; 1:200). The secondary antibodies were used at 1:1000 and all from Jackson ImmunoResearch: donkey anti-goat (705-545-147), donkey anti-mouse (705-585-151), donkey anti-rabbit (711-585-152, 711-545-152), donkey anti-rat (712-585-153), donkey anti-chicken (703-606-155).

For whole-mount X-gal staining, staged embryos were fixed with freshly-prepared, ice-cold 2% PFA and 0.2% glutaraldehyde in PBS for 1 hour at 4◻, rinsed with staining solution (5mM potassium ferricyanide, 5mM potassium ferrocyanide, 2mM MgCl_2_, 0.01% sodium deoxycholate, and 0.02% IGEPAL CA-630), followed by addition of X-gal stock solution (40mg/ml in dimethylformamide) at 1:40. For section X-gal staining, fixed embryos were dehydrated through gradient sucrose/OCT, embedded in OCT, cryo-sectioned at 10μm, and stained for X-gal as described above.

### Western Blot and real-time quantitative PCR (RT-qPCR)

P0 pups of wild type and *Gja5*^Δ14/Δ14^ mice were euthanized by decapitation. Hearts were then isolated, and atria were separated from ventricles. Samples from about 20 pups of each genotype were pooled, homogenized, and subjected to protein extraction or RNA extraction followed by cDNA preparation, as previously described (Li et al., 2019; Ye et al., 2015). For Western Blot, the following primary antibodies were used: anti-Cx40 (Cx40-A; Alpha Diagnostic International; 1:2000), anti-GAPDH (2118S; Cell Signaling Technology; 1:2000). The secondary antibody used was: IRDye® 800CW Donkey anti-Rabbit (926-32213; LI-COR; 1:10000). For RT-qPCR, the following primers were used: *Gja5* (F, 5’ -GGTCCACAAGCACTCCACAG-3’; R, 5′-CTGAATGGTATCGCACCGGAA-3′), *Gapdh* (F, 5’ -ATCAAGAAGGTGGTGAAGCAG-3’; R, 5′ -GAGTGGGAGTTGCTGTTGAAGT-3′).

### Chromatin Conformation Capture (3C) assays

Briefly, embryonic hearts were dissected under a microscope, minced into small pieces and digested with Accutase (Invitrogen 00-4555-56) at 37◻ for about 20 minutes, passed through a 70μm cell strainer, followed by formaldehyde fixation and quenched with glycine. The cells were then cryoprotected at −80◻. 1×10^7^ cells of each genotype (about 30 wild type embryos or 50 *Nkx2-5*^Cre/Cre^ embryos) were pooled for each 3C library preparation following standard procedures (Cope and Fraser, 2009). The relative interaction frequency (RIF) of each fragment of interest was measured by RT-qPCR, as described previously (Li et al., 2019). Primer details are as follows: F1 (5’ -GCCAAGGCCCTCAAGGTGA-3’), F2 (5’ -GGAGGGATTTGATATGAATGTAAGCACTG-3’), F3 (5’ -ATTCATGTAAGAGGGTCTGATCTCCAG-3’), F4 (5’ -ACAACCTTATCTCCAAACCTTTGCTC-3’), R1 (5’ -CCATTCCCTTTAGGACGGTTACCTTC-3’), R2 (5’ -CATGGACTGACCTCATTGGAGTG-3’), R3 (5’ -GGGAGGGATTCAGGACATGTTG-3’).

### Statistical analysis

All experiments were repeated at least three times to ensure scientific reproducibility. Quantification results are presented as mean±s.e.m., and statistical analysis was conducted using Student’s *t*-test in a GraphPad Prism 6 software. *P*<0.05 was considered significant.

## Acknowledgments

We thank Mrs. Ann Mullin at the Tulane Transgenic Animal Core Facility for assistance with the generation of genetically modified mice.

## Author contributions

Conceptualization: Y.C., T.Y.; Methodology: Y.C., T.Y., Z.H.; Validation: Y.C., T.Y.; Formal analysis: T.Y., Z.H., H.L.; Investigation: T.Y., Z.H., H.L., L.W.; Resources: Y.C.; Data curation: Y.C., T.Y.; Writing-original draft: T.Y.; Writing – review & editing: Y.C.; Visualization: T.Y.; Supervision: Y.C.; Project administration: Y.C.; Funding acquisition: Y.C.

