## Supplementary_File for "Conjugated activation of myocardial-specific transcription of *Gja5* by a pair of Nkx2-5-Shox2 co-responsive elements"

### Supplemental Figure Legends

#### Figure S1

**Figure S1. *Gja5-S1* and *Gja5-S2* are co-occupied by Shox2 and Nkx2-5 and marked as putative open chromatin.**
ChIP-seq datasets of Shox2 and Nkx2-5 on E12.5 embryonic atria are integrated with datasets of Assay for Transposase Accessible Chromatin with high-throughput sequencing (ATAC-seq) on (from top to bottom) E12.5 hearts, facial tissues, forebrains, hindbrains, livers, midbrains and neural tissues (ENCODE project). *Gja5-S1* and *Gja5-S2* are highlighted in light-blue vertical lines.

#### Figure S2

**Figure S2. *Gja5-eh* recapitulates endogenous *Gja5* expression in *Nkx2-5*$^+$ myocardium**
Triple immunofluorescent staining (Nkx2-5, Cx40, β-gal) on left ventricle (A-D), right ventricle (E-L) and right atrium (M-T) of the *Gja5-eh-LacZ* hearts at E12.5 and P60. CA, coronary arteries; VT, ventricular trabeculae; SAN, sinoatrial node. Scale bars: 100 μm.

#### Figure S3

**Figure S3. *Gja5-S2* is marked by H3K4me2, H3K4me3 and associated and occupied by CTCF.**
Integrative visualizing datasets of (from top to bottom) CTCF ChIP-seq on P0 hearts, H3K4me3 ChIP-seq on E13.5, E16.5, P0 hearts, and H3K4me2 ChIP-seq on E13.5, E16.5, P0 hearts. *Gja5-S1* and *Gja5-S2* are highlighted in light-blue vertical lines.

#### Figure S4

**Figure S4. Predicted transcription factor binding sites within *Gja5-S1***
Integrative visualizing datasets of (from top to bottom) Shox2 ChIP-seq, Nkx2-5 ChIP-seq on E12.5 hearts, and JASPAR-2020 transcription factor binding profiling on *Gja5-S1*. Predicted binding sites of Nkx2-3, Nkx2-5 and Nkx2-8 are highlighted in red rectangles.

#### Figure S5

**Figure S5. Predicted transcription factor binding sites within *Gja5-S2***
Integrative visualizing datasets of (from top to bottom) Shox2 ChIP-seq, Nkx2-5 ChIP-seq on E12.5 hearts, and JASPAR-2020 transcription factor binding profiling on *Gja5-S2*. Predicted binding sites of Nkx2-3 and Nkx2-8 are highlighted in red rectangles.

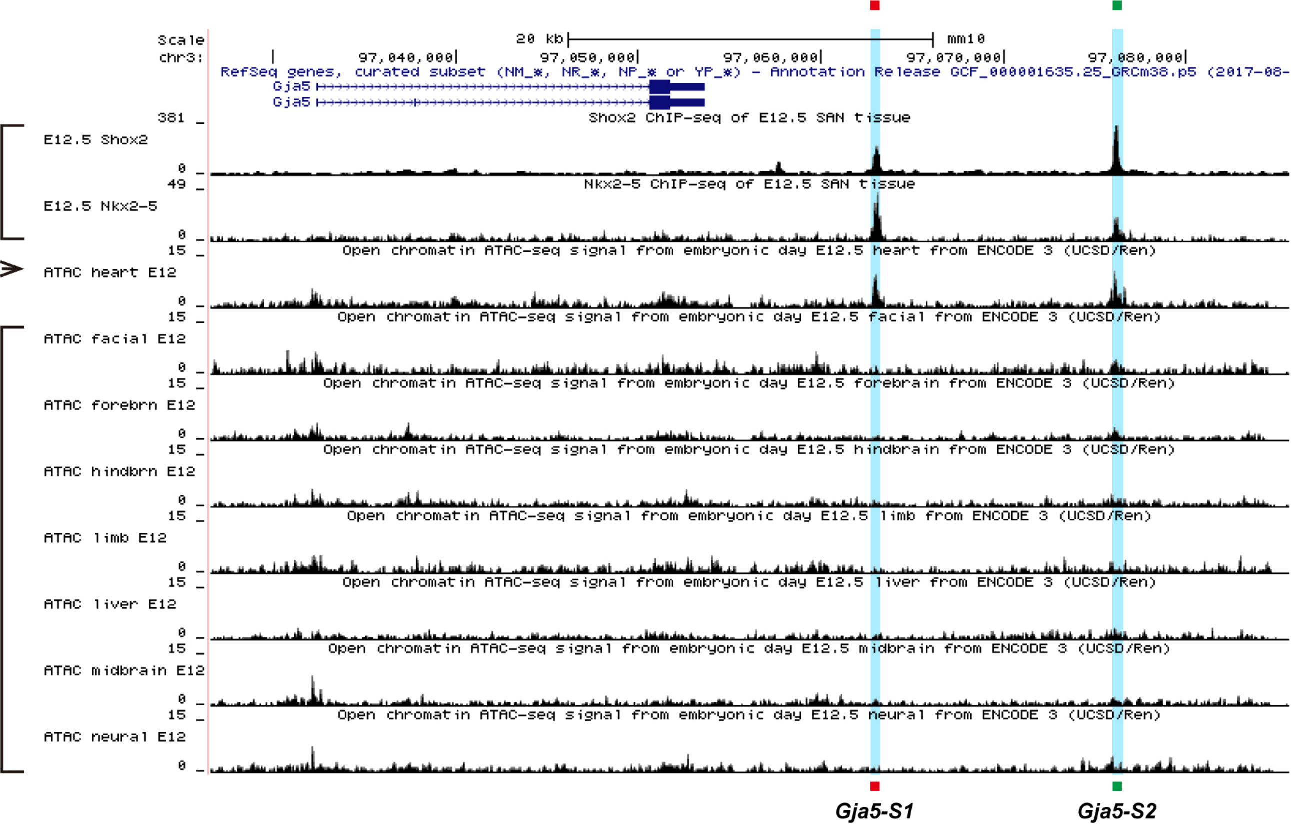

Scale
chr3:
20 kb
mm10
97,040,000
97,050,000
97,060,000
97,070,000
97,080,000
RefSeq genes, curated subset (NM_*, NR_*, NP_* or YP_*) - Annotation Release GCF_000001635.25_GRCm38.p5 (2017-08-
Gja5
Gja5
381
Shox2 ChIP-seq of E12.5 SAN tissue
E12.5 Shox2
0
49
Nkx2-5 ChIP-seq of E12.5 SAN tissue
E12.5 Nkx2-5
0
15
Open chromatin ATAC-seq signal from embryonic day E12.5 heart from ENCODE 3 (UCSD/Ren)
ATAC heart E12
0
15
Open chromatin ATAC-seq signal from embryonic day E12.5 facial from ENCODE 3 (UCSD/Ren)
ATAC facial E12
0
15
Open chromatin ATAC-seq signal from embryonic day E12.5 forebrain from ENCODE 3 (UCSD/Ren)
ATAC forebrn E12
0
15
Open chromatin ATAC-seq signal from embryonic day E12.5 hindbrain from ENCODE 3 (UCSD/Ren)
ATAC hindbrn E12
0
15
Open chromatin ATAC-seq signal from embryonic day E12.5 limb from ENCODE 3 (UCSD/Ren)
ATAC limb E12
0
15
Open chromatin ATAC-seq signal from embryonic day E12.5 liver from ENCODE 3 (UCSD/Ren)
ATAC liver E12
0
15
Open chromatin ATAC-seq signal from embryonic day E12.5 midbrain from ENCODE 3 (UCSD/Ren)
ATAC midbrain E12
0
15
Open chromatin ATAC-seq signal from embryonic day E12.5 neural from ENCODE 3 (UCSD/Ren)
ATAC neural E12
0
Gja5-S1
Gja5-S2

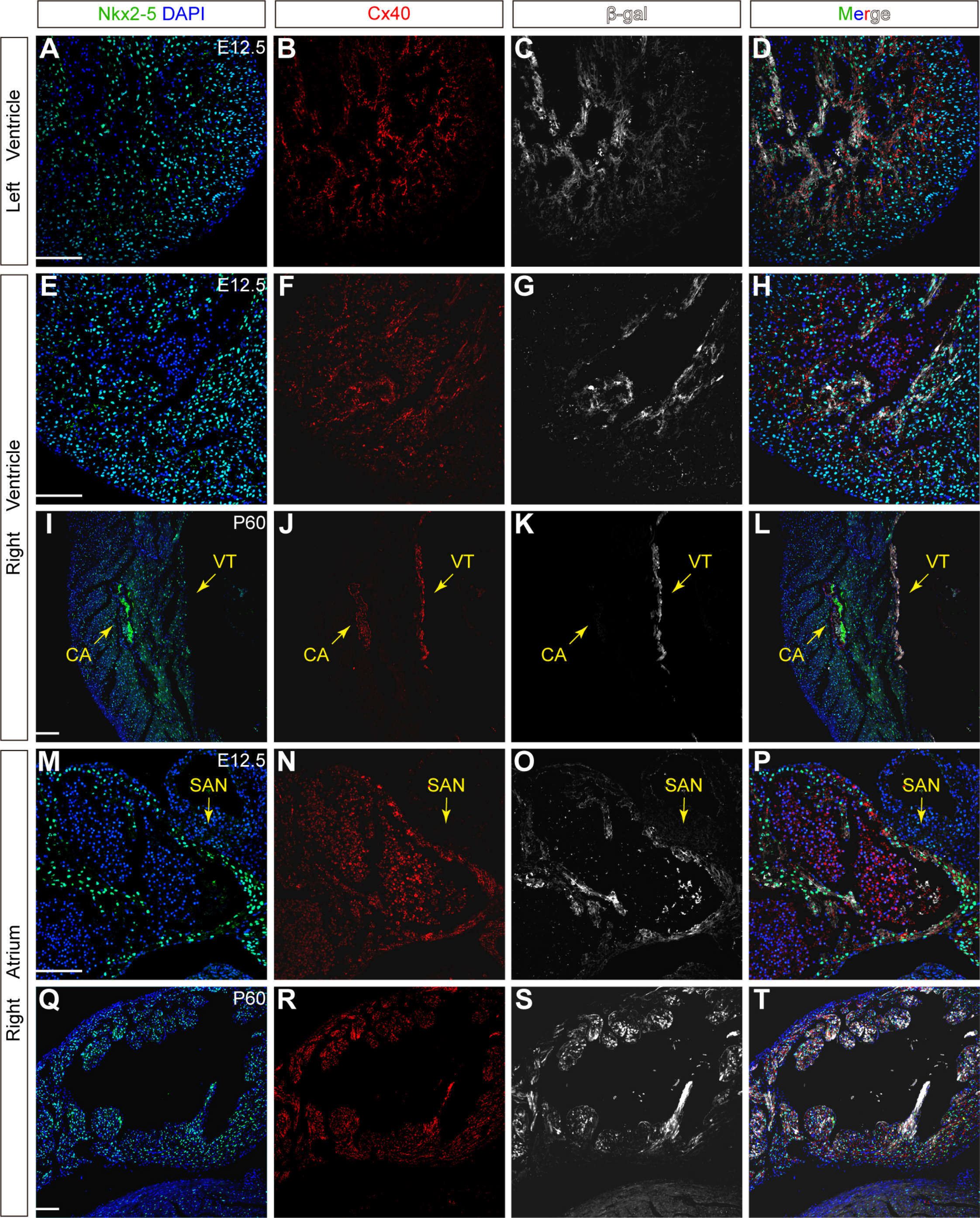

Nkx2-5 DAPI
Cx40
β-gal
Merge
Left Ventricle
Right Ventricle
Right Atrium
A
E12.5
B
C
D
E
E12.5
F
G
H
I
P60
VT
CA
J
VT
CA
K
VT
CA
L
VT
CA
M
E12.5
SAN
N
SAN
O
SAN
P
SAN
Q
P60
R
S
T

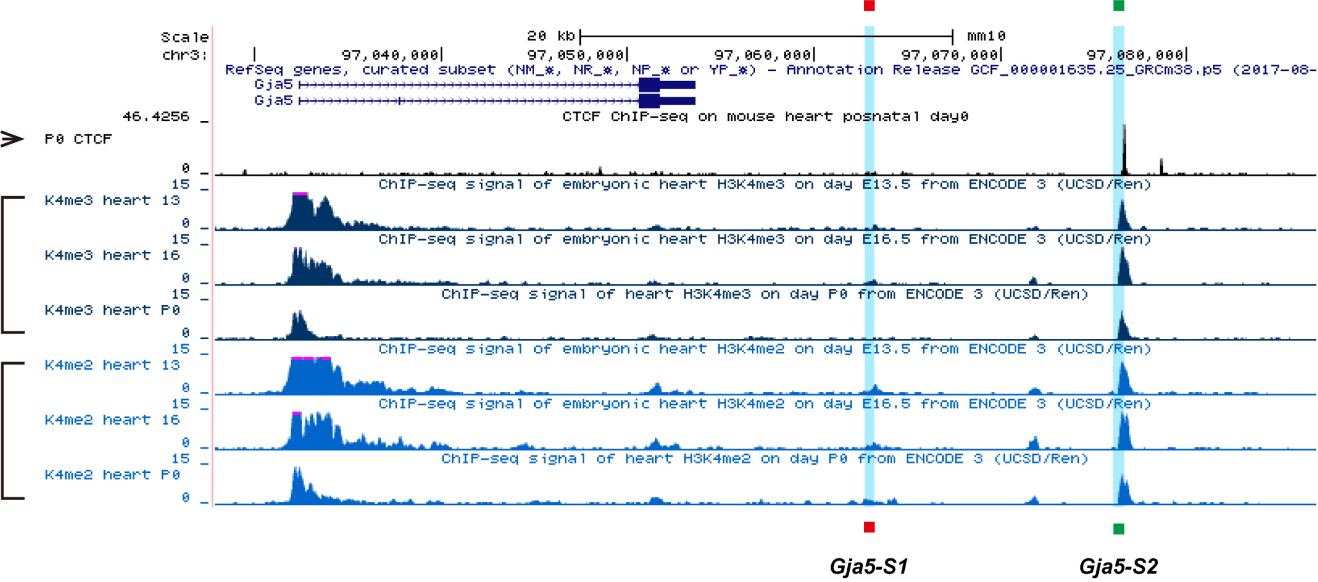

Scale
20 kb
mm10
chr3:
97,040,000
97,050,000
97,060,000
97,070,000
97,080,000
RefSeq genes, curated subset (NM_*, NR_*, NP_* or YP_*) - Annotation Release GCF_000001635.25_GRCm38.p5 (2017-08-
Gja5
Gja5
46.4256
CTCF ChIP-seq on mouse heart posnatal day0
P0 CTCF
0
15
ChIP-seq signal of embryonic heart H3K4me3 on day E13.5 from ENCODE 3 (UCSD/Ren)
K4me3 heart 13
0
15
ChIP-seq signal of embryonic heart H3K4me3 on day E16.5 from ENCODE 3 (UCSD/Ren)
K4me3 heart 16
0
15
ChIP-seq signal of heart H3K4me3 on day P0 from ENCODE 3 (UCSD/Ren)
K4me3 heart P0
0
15
ChIP-seq signal of embryonic heart H3K4me2 on day E13.5 from ENCODE 3 (UCSD/Ren)
K4me2 heart 13
0
15
ChIP-seq signal of embryonic heart H3K4me2 on day E16.5 from ENCODE 3 (UCSD/Ren)
K4me2 heart 16
0
15
ChIP-seq signal of heart H3K4me2 on day P0 from ENCODE 3 (UCSD/Ren)
K4me2 heart P0
0
Gja5-S1
Gja5-S2

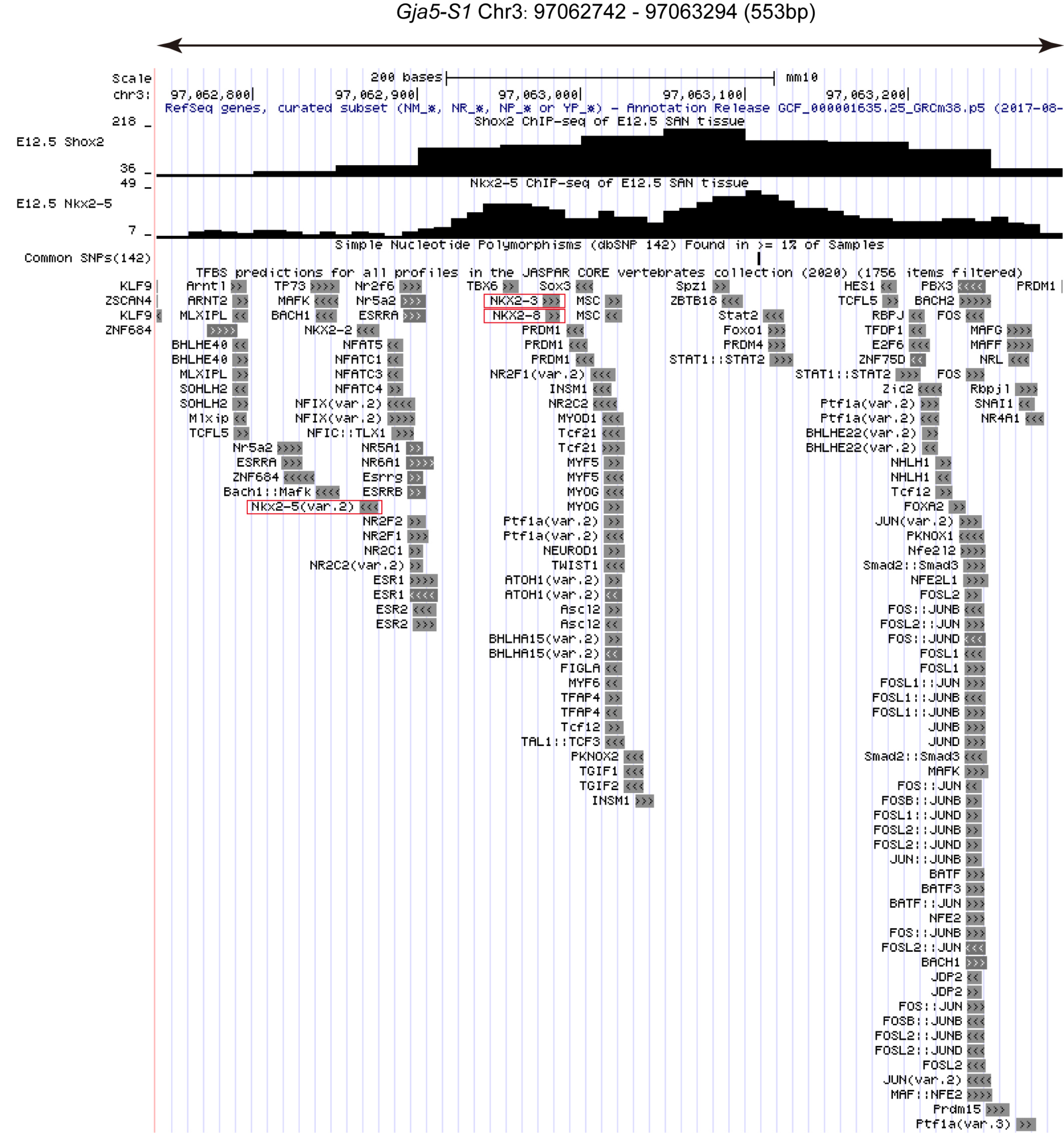

Gja5-S1 Chr3: 97062742 - 97063294 (553bp)
Scale
200 bases
mm10
chr3:
97,062,800
97,062,900
97,063,000
97,063,100
97,063,200
RefSeq genes, curated subset (NM_*, NR_*, NP_* or YP_*) - Annotation Release GCF_000001635.25_GRCm38.p5 (2017-08-
Shox2 ChIP-seq of E12.5 SAN tissue
218
E12.5 Shox2
36
49
Nkx2-5 ChIP-seq of E12.5 SAN tissue
E12.5 Nkx2-5
7
Simple Nucleotide Polymorphisms (dbSNP 142) Found in >= 1% of Samples
Common SNPs(142)
TFBS predictions for all profiles in the JASPAR CORE vertebrates collection (2020) (1756 items filtered)
KLF9
ZSCAN4
KLF9
ZNF684
Arntl
ARNT2
MLXIPL
BHLHE40
BHLHE40
MLXIPL
SOHLH2
SOHLH2
Mlxip
TCFL5
Nr5a2
ESRRA
ZNF684
Bach1::Mafk
Nkx2-5(var.2)
TP73
MAFK
BACH1
Nr2f6
Nr5a2
ESRRA
NKX2-2
NFAT5
NFATC1
NFATC3
NFATC4
NFIX(var.2)
NFIX(var.2)
NFIC::TLX1
NR5A1
NR6A1
Esrrg
ESRRB
NR2F2
NR2F1
NR2C1
NR2C2(var.2)
ESR1
ESR1
ESR2
ESR2
TBX6
Sox3
NKX2-3
NKX2-8
MSC
MSC
PRDM1
PRDM1
PRDM1
NR2F1(var.2)
INSM1
NR2C2
MYOD1
Tcf21
Tcf21
MYF5
MYF5
MYOG
MYOG
Ptf1a(var.2)
Ptf1a(var.2)
NEUROD1
TWIST1
ATOH1(var.2)
ATOH1(var.2)
Ascl2
Ascl2
BHLHA15(var.2)
BHLHA15(var.2)
FIGLA
MYF6
TFAP4
TFAP4
Tcf12
TAL1::TCF3
PKNOX2
TGIF1
TGIF2
INSM1
Spz1
ZBTB18
Stat2
Foxo1
PRDM4
STAT1::STAT2
HES1
TCFL5
RBPJ
TFDP1
E2F6
ZNF75D
STAT1::STAT2
Zic2
Ptf1a(var.2)
Ptf1a(var.2)
BHLHE22(var.2)
BHLHE22(var.2)
NHLH1
NHLH1
Tcf12
FOXA2
JUN(var.2)
PKNOX1
Nfe2l2
Smad2::Smad3
NFE2L1
FOSL2
FOS::JUNB
FOSL2::JUN
FOS::JUND
FOSL1
FOSL1
FOSL1::JUN
FOSL1::JUNB
FOSL1::JUNB
JUNB
JUND
Smad2::Smad3
MAFK
FOS::JUN
FOSB::JUNB
FOSL1::JUND
FOSL2::JUNB
FOSL2::JUND
JUN::JUNB
BATF
BATF3
BATF::JUN
NFE2
FOS::JUNB
FOSL2::JUN
BACH1
JDP2
JDP2
FOS::JUN
FOSB::JUNB
FOSL2::JUNB
FOSL2::JUND
FOSL2
JUN(var.2)
MAF::NFE2
Prdm15
Ptf1a(var.3)
PBX3
BACH2
FOS
MAFG
MAFF
NRL
FOS
Rbpjl
SNAI1
NR4A1
PRDM1

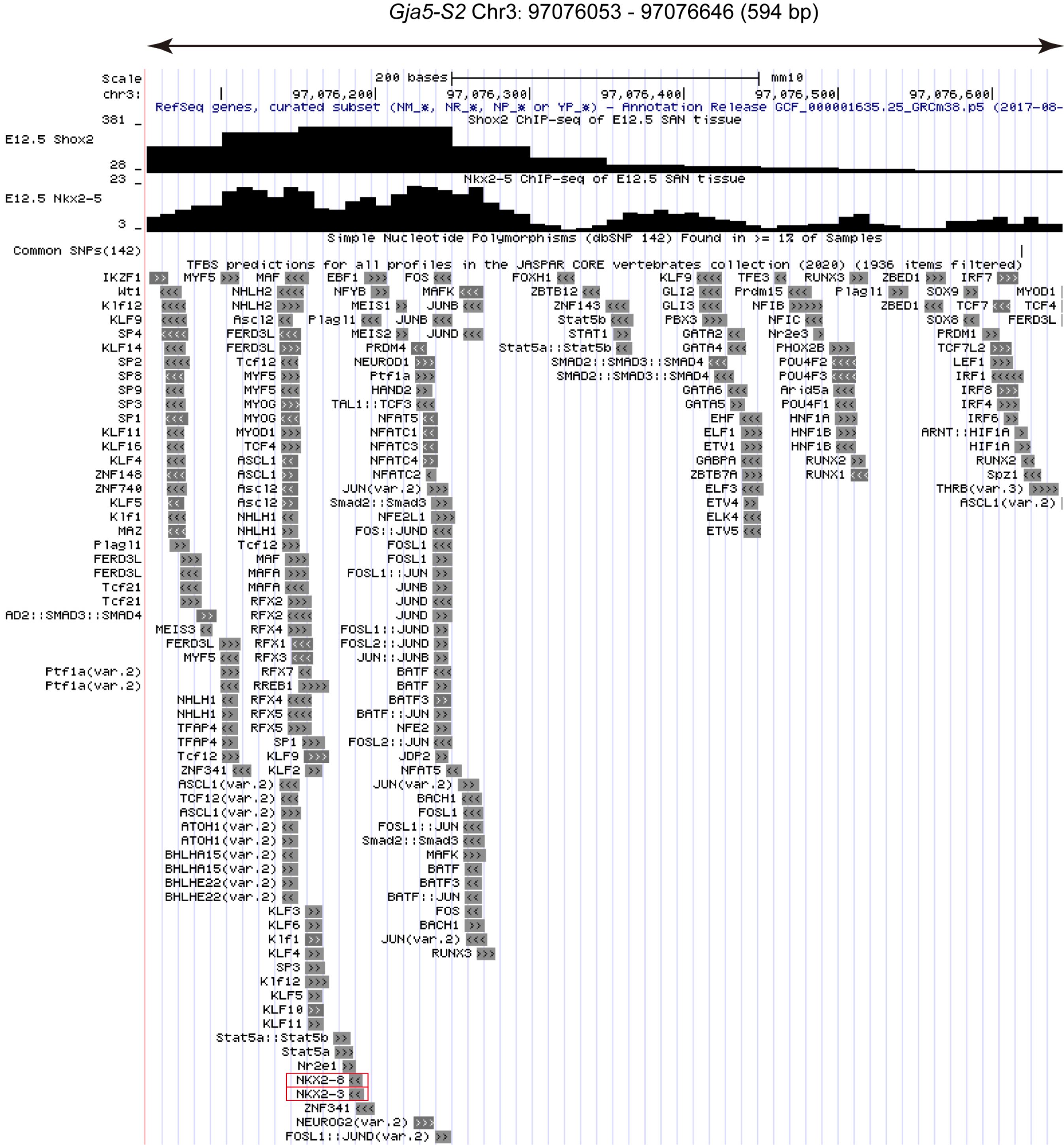
Gja5-S2 Chr3: 97076053 - 97076646 (594 bp)
Scale
200 bases
mm10
chr3:
97,076,200
97,076,300
97,076,400
97,076,500
97,076,600
RefSeq genes, curated subset (NM_*, NR_*, NP_* or YP_*) - Annotation Release GCF_000001635.25_GRCm38.p5 (2017-08-
381 _
Shox2 ChIP-seq of E12.5 SAN tissue
E12.5 Shox2
28 _
23 _
Nkx2-5 ChIP-seq of E12.5 SAN tissue
E12.5 Nkx2-5
3 _
Simple Nucleotide Polymorphisms (dbSNP 142) Found in >= 1% of Samples
Common SNPs(142)
TFBS predictions for all profiles in the JASPAR CORE vertebrates collection (2020) (1936 items filtered)
NKX2-8
NKX2-3

### Supplementary Materials and Methods

#### Generation and genotyping of transgenic mice

Generation and genotyping of transgenic mice were performed as described previously (Ye et al., 2016). Two permanent *Gja5-eh-LacZ* transgenic mouse lines were established by backcrossing with C57/BL6 wild type mice for at least six generations (over a period of 14 months). As we observed identical enhancer activity in both lines compared to that of F0 transgenic embryos (data not shown), we subsequently maintained one line for our studies. The primers of genotyping *Gja5-eh-LacZ* transgenic mouse lines are: Forward (5' -CCCATCTAGT CGCACTCTGG- 3') and Reverse (5' -GGAGAAGTGCAGAAAACCATGC- 3').

#### Generation and analysis of CRISPR/Cas9-mediated genetically modified mice

The $Gja5^{\Delta14}$ and $Gja5^{\Delta12}$ alleles were generated by CRISPR/Cas9-mediated genome editing (Wang et al., 2013). The sgRNAs were selected using an online platform (www.benchling.com) to optimize on-target/off-target potential (Doench et al., 2016; Hsu et al., 2013). For the $Gja5^{\Delta14}$ allele, the following pair of sgRNAs (Protospacer-adjacent motif (PAM) sequences are indicated in underlined bases) were used: sgRNA-1F (5' -ACTTTCTTTCACCATCTTGTGG G- 3') and sgRNA-1R (5' -TGGTGGTGCCACCTACAGATGGG- 3'). The sequence length between sgRNA-F1 and sgRNA-R1 is 13973-bp (~14 Kb). For the $Gja5^{\Delta12}$ allele, the following pair of sgRNAs were used: sgRNA-2F (5' -AAGTACCCTGCTACGTTGGTGGG- 3') and sgRNA-2R (5' -AAAGATCCCACGTTGTTGCGTGG- 3'). The sequence length between sgRNA-F2 and sgRNA-R2 is 12513-bp (~12 Kb). All predicted off-target sequences have at least three mismatches and located at least 10 Mb from the targeting sites. None of these potential off-targets were in significant linkage disequilibrium with *Gja5*.

The Cas9 proteins and sgRNAs were commercially provided (idtdna.com). Preparation of CRISPR injection mixes and microinjection of one-cell embryos were performed according to established protocols (Quadros et al., 2017). Multiple F0 founders bearing the predicted deletion were obtained, and two founders of each genotype were outcrossed to wild type

C57/BL6 mice to establish independent lines. F1 offsprings were then intercrossed to generate F2 mice where homozygous deletion of each allele was obtained. The presence of wild type and deleted alleles was confirmed by genotyping using the following primers: *Gja5*$^{\Delta14}$-F (5' -CACCCACTTATTGGGGCACTAT- 3') and *Gja5*$^{\Delta14}$-R (5' -CCCCAGAGGAGGTCAAACT G- 3') for genotyping of the *Gja5*$^{\Delta14}$ allele (expected PCR product: 405-bp); *Gja5*$^{\Delta12}$-F (5' -CCCTGGAAAGTCAGCACACC- 3') and *Gja5*$^{\Delta12}$-R (5' -CATCTGTTTCAACTCTGTCAA CACACAG- 3') for genotyping of the *Gja5*$^{\Delta12}$ allele (expected PCR product: 449-bp); *Gja5*-internal-F (5' -AGAGGTGATGCCAGGAAGGA- 3') and *Gja5*-internal-R (5' -GCATTGGA TCTGTCTGTTGCC- 3') for genotyping of the wild type allele with a 502-bp PCR product that is absent in either *Gja5*$^{\Delta14}$ or *Gja5*$^{\Delta12}$ allele.
